# Statistical Olfactory Learning in Honey Bees

**DOI:** 10.1101/2025.03.23.644064

**Authors:** Claire Marcout, Chiara Santolin, Martin Giurfa, Marco Paoli

## Abstract

Statistical learning is a key mechanism for detecting regularities in sensory inputs^1,2^. Among its functions is the ability to extract regularities from sequences (of sounds, objects, odors, etc.)^3^, enabling species to predict future events and guide behavior. This capacity has been demonstrated in vertebrates^2^, including human infants^4^, non-human primates^5^, and birds^6^. However, the minimum computational architecture required for statistical learning remains unclear. To address this issue, we studied statistical learning in the honey bee (*Apis mellifera*), an invertebrate model for learning studies^7^. We show that bees learn and recall the temporal structure of sequences of odorants, suggesting that statistical learning is a fundamental component of a conserved cognitive toolkit present even in invertebrates.

## Main Text

Using the proboscis extension reflex (PER) protocol^8^, we trained bees to discriminate between sequences of the same two odorants, A and B, presented in opposite orders: AB vs. BA (**Fig. 1A**). The rewarded/conditioned sequence (CS+) (e.g., AB) was paired with an appetitive sugar solution, while the reversed sequence (CS-) (e.g., BA) was unrewarded. To control for odor-related biases, AB and BA were alternated as the rewarded sequence (CS+). Additionally, two odorant sets were used as A and B - i.e., 2-hexanol and nonanal (*n* = 223 bees) and 1-hexanol and 2-nonanol (*n* = 202 bees) - selected based on their perceptual differences in bees^9^ (see STAR Methods). For clarity, we will refer to AB as the CS+ and BA as the CS-.

**Figure 1.**
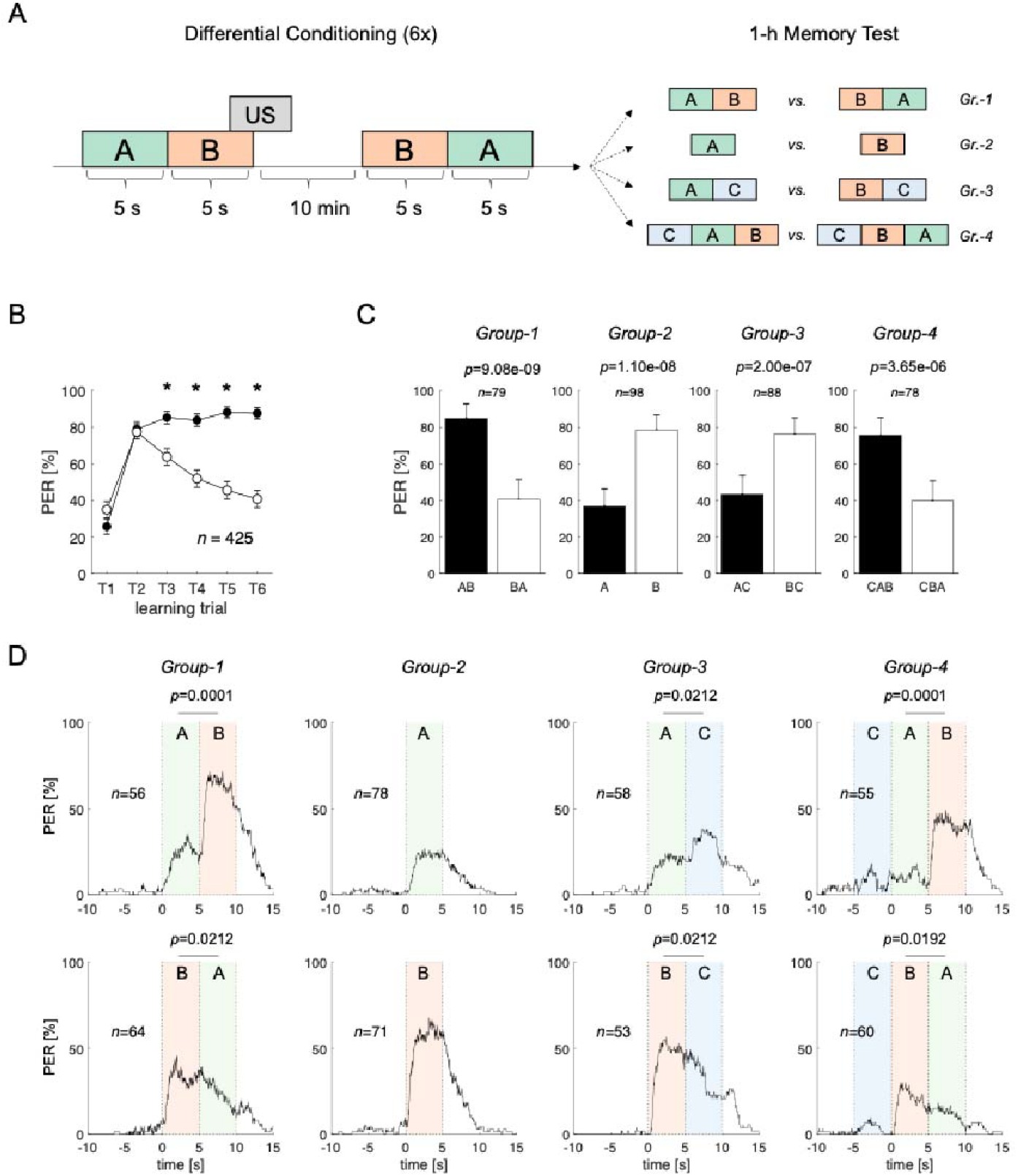
Differential sequence learning and 1-hour memory test in honey bees. **(A)** Scheme of the differential appetitive conditioning protocol: six exposures to the conditioned sequence (CS+, e.g., AB), each followed by a 3-s sucrose reward with a 1-s overlap with the second element of the sequence, and six exposures to the unrewarded reversed sequence (CS-, e.g., BA). One hour after the last learning trial, bees were divided into four groups and subjected to different memory retention tests. Abbreviations: Groups 1-4, Gr. 1-4; unconditioned stimulus, US. **(B)** Learning curve for the differential conditioning phase (*n*=425). Data represent the % of bees showing PER during the 10-s odor sequence, with 95% confidence intervals. GLMM analysis shows a significant response difference from trial three (T3) onward (*, *p*∼0). Abbreviation: proboscis extension response, PER. **(C)** Each conditioned bee underwent two 1-h memory tests to evaluate the relevance of different features of the conditioned stimulus for memory recall. Data represent the % of bees showing PER during the test sequence, with 95% confidence intervals. The number of individuals (*n*) and *p*-values from the McNemar’s test comparing response rates to the tested stimuli are displayed above each plot. **(D)** Memory tests were video recorded and proboscis extension was tracked to quantify response dynamics during olfactory stimulation. Differences in proboscis responses probabilities [%] to the different stimuli within each pattern were tested using a Wilcoxon signed-rank test. *P*-values corrected for false discovery rate (FDR) are shown above each plot.

For both sets of odorants, bees discriminated the sequences from the third training trial onwards (GLMM, trial/stimulus interaction effect, from T3 to T6 *p*∼0.000; **Fig. 1B**). Discrimination improved across trials, indicating progressive task learning. Since both odorants predicted the presence or absence of food reward, recognition of the correct sequence must have relied on the relative positions of the odorants. One hour later, the conditioned bees were divided into four groups for memory tests to investigate the sequence features relevant to statistical learning.

*Group-1* bees were presented with the conditioned and reversed odorant sequences (i.e., AB and BA) and showed significant recognition of the CS+ and the CS-(McNemar’s test, *n*=79, *p*=9.1*10^−9^; **Fig. 1C**). This indicates that the previously acquired information on the odorant sequences was retained for at least one hour.

*Group-2* bees were tested with individual odorants (i.e., only A or B) to assess whether they memorized the sequences as whole entities or whether they focused on each individual odorant. Bees showed a significantly higher PER probability to the second odorant of the previously rewarded sequence (*n*=98, *p*=1.1*10^−8^; **Fig. 1C**). This suggests that, in addition to discriminating odorant sequences based on the relative positions of the odorants, bees responded significantly more to the second element, assigning it greater weight due to its proximity to the reward in half of the learning trials.

*Group-3* bees were tested with novel combinations, in which the second odorant of rewarded and unrewarded sequences was replaced by a novel, neutral odorant C, well-perceived by bees^9^ (i.e., A**B**→A**C**; B**A**→B**C**). Bees trained with the first odorant set (2-hexanol/nonanal) received 2-octanol as the novel odorant C, while bees trained with the second set (1-hexanol/2-nonanol), received 2-hexanone. This manipulation aimed to test whether the first odorant acquired predictive value, leading to higher PER to C in AC but not in BC. However, bees responded significantly more to BC, which included B, the odorant adjacent to the reward during training (*n*=88, *p*=`2.0*10^−7^; **Fig. 1C**).

*Group-4* bees were tested with two three-odorant sequences, where the novel odorant C was added at the beginning of rewarded and unrewarded sequences (i.e., **C**AB and **C**BA, respectively). This test assessed whether odorants’ positional information served as a cue for memory retrieval. During differential conditioning with AB (CS+) and BA (CS-), B was rewarded when it occurred in the second position. If positional information was critical, bees should show higher PER probability to CBA, where B remains in the second position. Conversely, if bees relied on the whole structure of the conditioned sequence, they should favor CAB, which comprises AB (the rewarded sequence). Bees conditioned with AB responded significantly more to CAB than to CBA (*n*=78, *p*=3.6*10^−6^), indicating that they encoded the whole AB sequence rather than just the position of individual odorants (**Fig. 1C**).

All memory tests were video-recorded and analyzed to track PER dynamics during olfactory stimulation (**Fig. 1D**). The mean response probability across bees was higher when the second element of the rewarded sequence (i.e., B) appeared, suggesting that bees relied on it as a primary cue during memory retrieval. Nevertheless, bees in *Group-3*, tested with AC and BC, significantly increased their appetitive response for the novel odorant C when it followed A (*n*=58, *p*=0.021) but decreased it when it followed B (*n*=53, *p*=0.021). This result indicates that A acquired appetitive predictive appetitive value. Notably, the increased response did not occur after exposure to A alone (**Fig. 1D**, *Group-2*) nor was it triggered by C alone, as shown by the low responsiveness to C when presented at the sequence onset (**Fig. 1D**, *Group-4*).

Our findings demonstrate that bees can encode temporal relationships between sequentially organized odorants and use this information to guide their behavior. They learn and recall a rewarded odor sequence, either alone or embedded within a more complex olfactory context (e.g., three-element sequence). Furthermore, they assign greater importance to the odorant closest to reward. Finally, the increased response to a neutral odorant following the first odorant of the rewarded sequence (**Fig 1D**, *Group-3*) highlights the bees’ sensitivity to the temporal structure of the olfactory input.

Statistical learning enables learning of temporal regularities across sensory modalities such as vision, audition, and haptics in vertebrate species with different evolutionary histories^1,2^. In humans and songbirds, it supports higher-order functions like early language acquisition and song learning^2,10^. Here, we showed that an invertebrate species - the honey bee - can learn temporal regularities defining olfactory sequences and detect changes in those patterns. This ability may assist bees in spatial orientation and foraging by signaling the proximity of landmarks or food sources, offering an evolutionary advantage for navigating complex olfactory environments.

## Supporting information

STAR Methods

## Acknowledgments

C.M, M.P. and M.G. thank the CNRS, INSERM and Sorbonne University for support. M.G was supported by an ERC Advanced Grant (COGNIBRAINS, 835032) and by the Institut Universitaire de France (IUF). C.S. was supported by an ERC Starting grant ‘GALA’ (101115991).

## Author contributions

Conceptualization, M.P., M.G., and C.S.; investigation, C.M. and M.P.; methodology, C.M., M.P., M.G. and C.S.; data curation, M.P.; visualization, M.P.; Software, M.P.; supervision, M.P., M.G., and C.S.; project administration, M.P., M.G., and C.S.; funding acquisition, M.G.; writing – original draft, M.P. and C.M.; writing – review and editing, C.M., M.P., M.G. and C.S..

## Declaration of interests

The authors declare no competing interests.

## Supplementary Figures

**Supplementary Figure 1.**
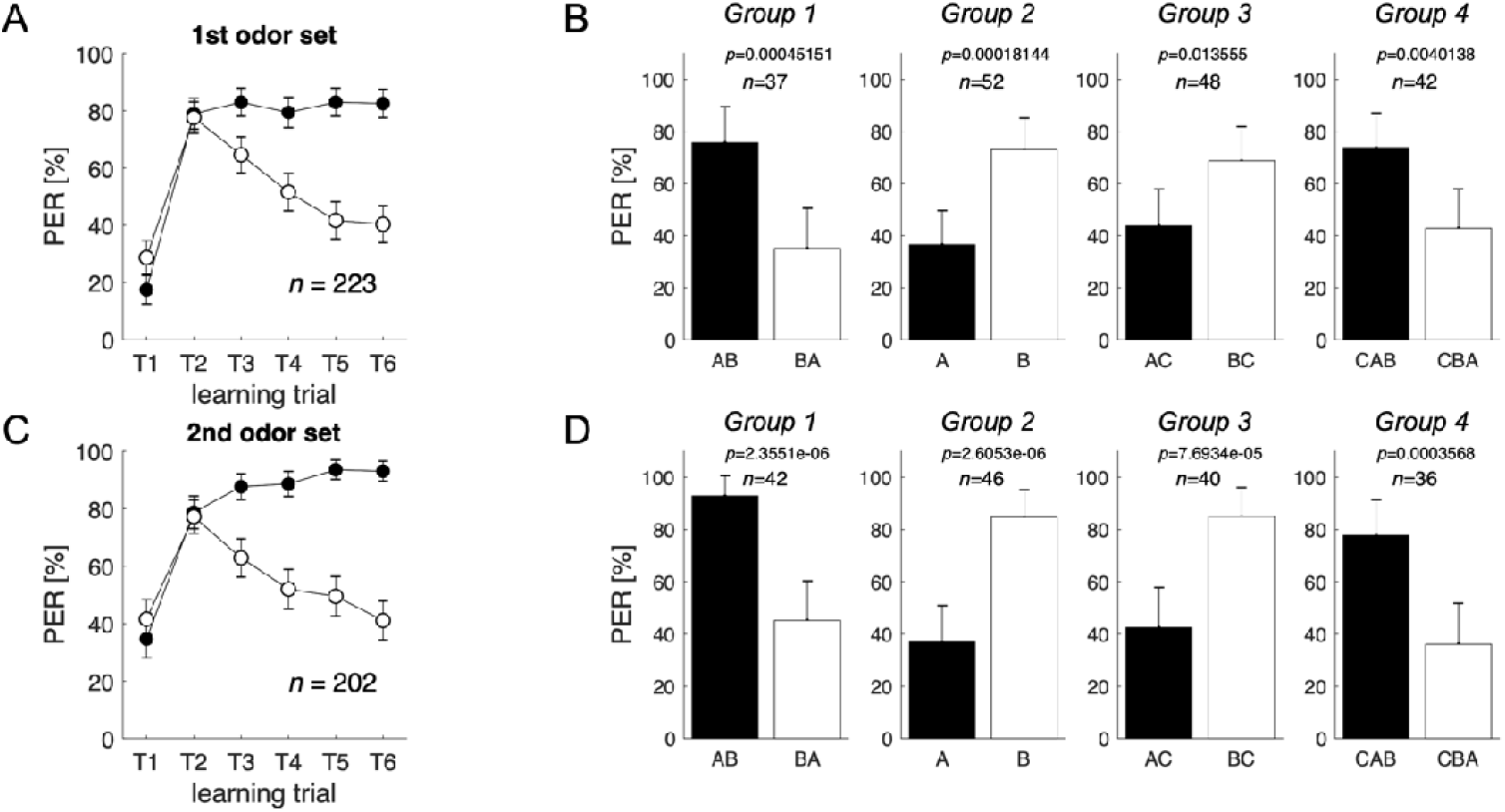
Differential sequence learning and 1-hour memory test for the two different odorant sets employed in the study. **(A)** Learning curve for the differential conditioning phase (*n*=223) for bees trained with 2-hexanol and nonanal (as A and B or vice versa). Data represent the % of bees showing PER during the 10-s sequence, with 95% confidence intervals. Abbreviation: proboscis extension response, PER. **(B)** Each conditioned bee from (**A**) underwent two 1-h memory tests to evaluate the relevance of different features of the conditioned stimulus for memory recall. Data represent the % of bees showing PER during the test sequence, with 95% confidence intervals. The number of individuals (*n*) and *p*-values from the McNemar’s test comparing response rates to the tested stimuli are displayed above each plot. **(C)** Learning curve for the differential conditioning phase (*n*=202) for bees trained with 1-hexanol and 2-nonanol (as A and B or vice versa). Data represent the % of bees showing PER during the 10-s sequence, with 95% confidence intervals. **(D)** Each conditioned bee from (**C**) underwent two 1-h memory tests to evaluate the relevance of different features of the conditioned stimulus for memory recall. Data represent the % of bees showing PER during the test sequence, with 95% confidence intervals. The number of individuals (*n*) and *p*-values from the McNemar’s test comparing response rates to the tested stimuli are displayed above each plot.

## Notes

https://github.com/mp599/2025_honeybee_statistical_learning

### Competing Interest Statement

The authors have declared no competing interest.

