## Supplementary material for "Statistical Olfactory Learning in Honey Bees": STAR Methods

### RESOURCES AVAILABILITY

#### Lead contact

Further information and requests should be directed to the lead contacts Chiara Santolin, Martin Giurfa and Marco Paoli, who will handle and fulfill these inquiries.

#### Materials availability

This study did not generate new unique reagents.

#### Data and code availability

All data and original code have been deposited on GitHub (https://github.com/mp599/2025_honeybee_statistical_learning) and are publicly available.

### EXPERIMENTAL MODEL AND SUBJECT DETAILS

#### Insects

Honey bee foragers from colonies of the Neuro-SU laboratory, located on the Pierre et Marie Curie Campus (Sorbonne Université, 75005, Paris), were collected in the afternoon and housed in custom-printed plastic cages in groups of 16 individuals. Each group was provided with 240 microliters of 50% sugar/water solution (an average of 15 μl/individual) and maintained overnight in a dark and humid incubator at 30°C. The following morning, the bees were individually harnessed in plastic tubes, secured with tape, and fed 3 μl of sugar solution. After a three-hour rest period, they underwent the olfactory conditioning protocol[1].

### METHOD DETAILS

#### Differential appetitive conditioning

Honey bees were subject to a differential conditioning protocol in which one odorant sequence (e.g., AB) was paired with a sugar reward, while the reversed sequence (e.g., BA) was not. Two sets of odorants were used for this experiment. One group of bees ($n$=223) was trained to discriminate between sequences of 2-hexanol (CAS number 626-93-7) and nonanal (CAS number 124-19-6). Half of these bees were trained with 2-hexanol followed by nonanal as the conditioned sequence (CS+), and with the reversed sequence (nonanal followed by 2-hexanol) as the non-conditioned stimulus (CS-). The remaining bees were trained with the opposite sequence assignments. A second group of bees ($n$=202) was trained with sequences composed of hexanol (CAS number 111-27-3) and 2-nonanol (CAS number 628-99-9). Half were trained with hexanol followed by 2-nonanol as CS+, while the rest were trained with the opposite sequence assignments. For the olfactory stimulation, undiluted odorants (1 mL) were placed in 20 mL glass vials and the headspace was delivered to the bees using an Arduino Uno-controlled odorant delivery device as previously described[2,3].

In the differential conditioning protocol, harnessed bees were positioned in front of the odorant delivery device. Each trial began with a 20-second exposure to odorless air (familiarization phase) followed by a 5-second presentation of the first odorant, a 5-second presentation of the second odorant, and a final 20-second exposure to odorless air. For the conditioned stimulus (CS+), a 50% sugar reward was provided for 3 seconds, beginning during the last second of the second odorant presentation. Each bee underwent six rewarded (CS+) and six unrewarded (CS-) trials presented in a pseudorandom order and with a 10-minute inter-trial interval.

#### Memory retention test

One hour after the last conditioning trial, bees were tested for memory retention using one of the following configurations of olfactory stimuli: (1) the conditioned sequences (e.g., AB and BA; *Group-1*); (2) the elemental components of the sequence (e.g., A and B; *Group-2*); (3) a novel sequence composed of the first element of the conditioned sequences followed by a novel odorant (e.g., AC and BC; *Group-3*); (4) the conditioned sequences preceded by a novel odorant (e.g., CAB and CBA; *Group-4*).

The tests occurred 10 minutes apart in a pseudorandom order. For bees conditioned with the first odorant set (i.e., 2-hexanol and nonanal), 2-octanol (CAS number 123-96-6) was adopted as the novel odorant (C). For bees conditioned with the second odorant set (i.e., 1-hexanol and 2-nonanol), 2-hexanone (CAS number 591-78-6) served as the novel odorant. All odorants were selected because of their perceptual difference for honey bees[4] and were purchased from Sigma-Aldrich (St. Louis, MO, USA).

### QUANTIFICATION AND STATISTICAL ANALYSIS

#### Video recording and tracking of proboscis extension

Memory retention tests were recorded at 50 fps using a Sony HandyCam HDR-CX625 video camera. Proboscis extension was tracked using the DeepLabCut software[5]. A total of 400 frames were extracted from videos of 20 individuals (20 frames/video) and manually labelled to generate the training dataset. For this study, the base, midpoint, and terminal tip of the proboscis were tracked when visible in the frame). After the first network training session (ResNet50, 500000 iterations), frames with low-likelihood labels were manually corrected to refine the model. The refined network was then used to track proboscis movement across all videos.

#### Data analysis

Data analysis was conducted using MatLab R2024b (The MathWorks Inc.). The trained network reliably identified the proboscis tip during extension, while its absence was consistently associated with low likelihood scores. Hence, a likelihood threshold of 0.8 was applied to detect proboscis extension. Bees that exhibited proboscis extension before stimulus onset were excluded from the analysis, as these instances prevented meaningful analysis of the relationship between olfactory stimulation and appetitive learning. Overall, 579 videos were acquired, of which 84 were discarded because of proboscis movement before stimulus onset, leaving 495 videos for analysis.

#### Statistical tests

Learning curves (**Fig. 1B**) were analyzed using a generalized linear mixed model (GLMM) using the *fitgmle* MatLab function. Trial, stimulus, and individual were included as categorical variables. The model assessed the effects of trial and stimulus, as well as their interaction (trial:stimulus), to evaluate learning progression and stimulus-dependent differences. A McNemar's test was used to compare proboscis extension response scores for the two olfactory sequences presented during the 1-hour memory retention test (**Fig. 1C**). A Wilcoxon signed-rank test was used to compare the average proboscis response activity between the first and second elements of the olfactory sequence. For each odorant, the mean proboscis response across individuals was calculated in the middle of the olfactory stimulation, i.e., between 2 and 3 seconds after odorant onset (**Fig. 1D**). A *post-hoc* false discovery rate (FDR) correction was applied to the resulting *p*-values.
